# TMEM2-mediated hyaluronan turnover maintains articular cartilage homeostasis during osteoarthritis progression

**DOI:** 10.64898/2025.12.09.693317

**Authors:** Tomoya Murotani, Toshihiro Inubushi, Yu Usami, Taisei Tomohiro, Wu Deyang, Shinnosuke Kusano, Renshiro Kani, Shamma Hisham, Yuki Shiraishi, Hiroshi Kurosaka, Fumitoshi Irie, Yu Yamaguchi, Takashi Yamashiro

**Author notes:** **Correspondence author:** Toshihiro Inubushi, Department of Orthodontics and Dentofacial Orthopedics, The University of Osaka Graduate School of Dentistry, Osaka 565-0871, Japan.

## Abstract

Osteoarthritis (OA) is a degenerative joint disease characterized by progressive disruption of the cartilage extracellular matrix (ECM), yet the molecular mechanisms governing ECM turnover during disease initiation remain incompletely defined. Hyaluronan (HA) is a major structural component of articular cartilage, and its regulated turnover is essential for maintaining tissue integrity. Transmembrane protein 2 (TMEM2) is a cell-surface hyaluronidase capable of degrading high–molecular weight HA under physiological conditions, but its role in joint tissues has remained unclear.

Here, we examine the spatiotemporal expression and functional contribution of TMEM2 in articular cartilage using single-cell transcriptomic analysis, histological approaches, and a chondrocyte-specific conditional knockout mouse model. Under physiological conditions, Tmem2 was predominantly expressed in non-calcified articular chondrocytes. Following joint destabilization, Tmem2 expression was transiently increased during early osteoarthritis, coinciding with reduced cartilage HA content, consistent with altered HA turnover. Importantly, genetic ablation of Tmem2 in chondrocytes markedly exacerbated osteoarthritis progression, resulting in accelerated cartilage delamination, increased chondrocyte apoptosis, reduced proliferative activity, and enhanced hypertrophic differentiation. These changes occurred without detectable abnormalities in the synovium, subchondral bone, or osteophyte formation, indicating a cartilage-intrinsic phenotype.

Collectively, these findings identify TMEM2 as an important regulator of hyaluronan homeostasis within the cartilage ECM and provide in vivo genetic evidence that TMEM2- dependent HA turnover contributes to the maintenance of articular cartilage integrity during osteoarthritis progression.

## Introduction

Osteoarthritis (OA) is a chronic degenerative joint disease characterized by the progressive breakdown of articular cartilage and secondary change in subchondral bone, synovium, and surrounding tissues^1^. As one of the most prevalent musculoskeletal disorders, OA is a major cause of disability, particularly in aging populations^2,3^. Currently, OA affects over 300 million people worldwide, and with increasing life expectancy, the number of affected individuals is projected to rise substantially by 2040^4,5^. Beyond its physical impact, OA impairs mobility, reduces quality of life, and imposes substantial economic burdens^6^.

The pathogenesis of OA is multifactorial, involving aging, mechanical stress, genetic predisposition, obesity, and chronic inflammation^7^. Yet the sequence of early molecular events that trriger matrix failure in cartilage and synovium remain unclear, and no curative treatment exists. Current strategies are largely symptomatic (lifestyle modification and analgesics), and intra-articular hyaluronan (HA) injections are used in some settings because HA depletion is common in OA joints⁸, although their long-term efficacy remains debated. These gaps underscore the need to define actionable mechanisms operating at disease onset.

HA is a high-molecular-weight glycosaminoglycan composed of repeating disaccharides of glucuronic acid and N-acetylglucosamine, often exceeding 10⁶ Da in size^8^. Abundant in synovial fluid and cartilage, HA plays essential roles in joint lubrication, water retention, and immune modulation^9–14^. It maintains the viscoelastic properties of synovial fluid, facilitates smooth joint movement, and contributes to cartilage integrity by regulating cellular processes such as chondrocyte proliferation and matrix homeostasis. HA homeostasis is tightly regulated through a balance of synthesis and degradation. The HAS (HA synthase) family enzymes, particularly HAS2, are responsible for HA biosynthesis, and their disruption leads to severe cartilage defects and abnormal joint development^15,16^. In OA, early pathological changes include the degradation of the HA-aggrecan network, which compromises cartilage integrity and accelerates disease progression^17^. A distinguishing feature of HA is its rapid turnover, with nearly one-third of the body’s HA (approximately 15 g in a 70 kg adult) undergoing degradation and renewal daily^18^. This dynamic metabolism occurs through an extracellular degradation process, generating intermediate HA fragments (10–100 kDa), followed by intracellular catabolism within lysosomes by hyaluronidases and exoglycosidases^19^. While HYAL1 and HYAL2, members of the HYAL family of hyaluronidases, were once thought to be the primary mediators of extracellular HA degradation, current evidence suggests their enzymatic activity is largely confined to intracellular compartments, where they function optimally at acidic pH^20–22^. This leaves a mechanistic gap regarding extracellular HA turnover at near-neutral pH within joint tissues.

TMEM2 (Transmembrane Protein 2), also known as CEMIP2, was initially identified as a large type II transmembrane protein with an unclear function. Studies in zebrafish tmem2 mutants revealed developmental defects, including endocardial cushion abnormalities and excessive accumulation of extracellular HA, suggesting a role for TMEM2 in HA metabolism^23,24^. Subsequent investigations demonstrated that TMEM2 functions as a cell-surface hyaluronidase, capable of degrading high-molecular-weight HA into smaller fragments at near-neutral pH, positioning it as a critical regulator of extracellular HA turnover^25^. The physiological significance of TMEM2 has been further underscored by findings in conditional *Tmem2* knockout mice, which exhibit abnormal craniofacial morphology and excessive HA accumulation, supporting its role as a major extracellular HA-degrading enzyme *in vivo*^26,27^. However, its role in joint tissues and contribution to OA pathogenesis remain unclear.

Recent work on joint HA catabolism has yielded mixed conclusions regarding TMEM2/CEMIP2. In synovial fibroblasts, TMEM2-dependent HA degradation and cytokine responsiveness appear limited, whereas CEMIP/HYBID is IL-6–inducible ^28,29^. In late-stage human OA cartilage, TMEM2 expression is not consistently elevated, suggesting stage dependence (Shiozawa 2022). Importantly, earlier discrepancies about intrinsic TMEM2 hyaluronidase activity, initially negative in some human assays^29,30^, have been largely resolved by assay optimization, substrate quality control, and inhibitor removal, with purified human and mouse TMEM2 now showing robust degradation of native HMW-HA, at levels comparable to HYAL2^31^. With the enzymology clarified, a central gap remains: does TMEM2 drive extracellular HA turnover *in vivo*, and if so, when and where within the joint does it act? To address this knowledge gap, we combined single-cell transcriptomic reanalysis with genetic and histological approaches in a post-traumatic osteoarthritis model to define the spatiotemporal distribution and physiological function of TMEM2 in articular cartilage. By mapping TMEM2 expression dynamics and determining the consequences of its conditional deletion in chondrocytes, we provide in vivo evidence that TMEM2 contributes to extracellular HA homeostasis and is required for the maintenance of cartilage integrity under joint stress.

## Results

### Tmem2 Expression Patterns in the Mouse Knee Joint

To gain insight into the role of Tmem2 in the knee joint, we reanalyzed publicly available single-cell RNA sequencing (scRNA-seq) data from normal mouse knee joints. Eight distinct cell clusters were identified. Among these, clusters 0, 2, 3, and 4 expressed chondrocyte markers (*Sox9*, *Col2a1*, and *Acan*) and were classified as chondrocytes (Figure 1A, D).

**Figure 1.**
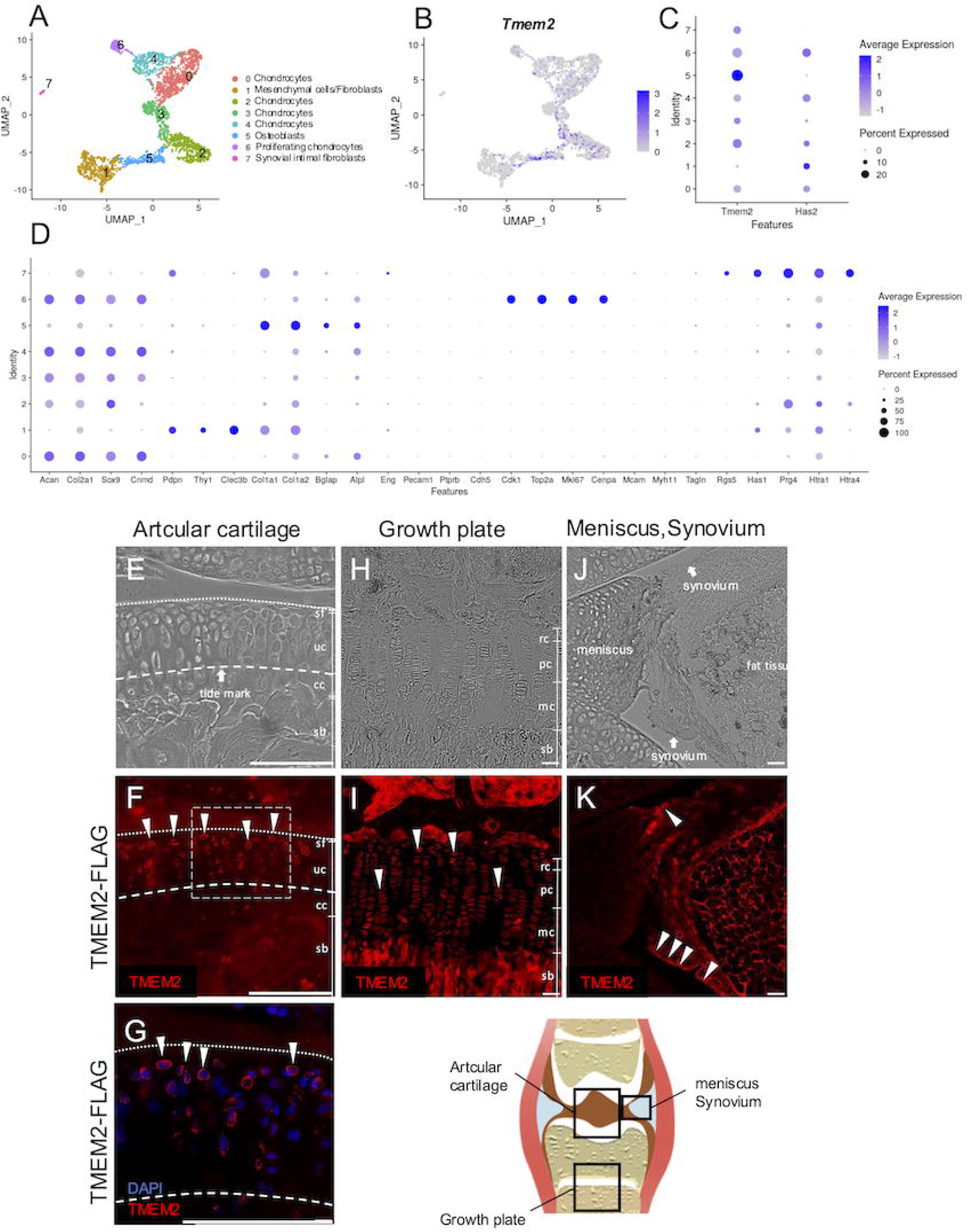
Localization of Tmem2 in the normal mouse knee joint and single-cell transcriptomic analysis of joint chondrocytes. (A) UMAP plot illustrating the distribution of 10 transcriptionally distinct cell populations identified in mouse knee joint chondrocytes, annotated by cell type. (B) UMAP feature plot showing the expression levels of *Tmem2* across cell clusters. The color gradient from grey to blue represents increasing gene expression intensity. (C) Dot plot comparing expression levels of *Tmem2* and *Has2* across clusters. *Tmem2* was highly expressed in chondrocytes (clusters 0, 2, 3, and 4), osteoblasts (cluster 5), proliferating chondrocytes (cluster 6), and synovial cells (cluster 7). (D) Dot plot presenting the expression of representative marker genes for each cell type, enabling cluster annotation. (E–F) Immunofluorescence images of sagittal frozen sections from the articular cartilage of 14-week-old Tmem2-FLAG mice. Tmem2 protein signal was enriched in the non-calcified cartilage layer (arrowheads). sf, superficial layer; uc, uncalcified cartilage; cc, calcified cartilage; sb, subchondral bone. (G) Higher-magnification image demonstrating Tmem2 protein localization on the chondrocyte plasma membrane (arrowheads). (H–I) Immunofluorescence analysis of the growth plate cartilage in 14-week-old Tmem2-FLAG mice. Strong Tmem2 protein expression was observed spanning the proliferative to hypertrophic zones of the growth plate (arrowheads). rc, resting cartilage; pc, proliferative cartilage; mc, mature cartilage; sb, subchondral bone. (J–K) Immunofluorescence images of the synovium and meniscus. Tmem2 protein was detected in synovial cells (arrowheads), which are involved in hyaluronic acid production, but not in the meniscal tissue. This experiment was performed with biological replicates (*n* = 5), and representative images are shown. Data shown in this figure are representative of at least three independent experiments. Scale bars: 20 μm.

Additionally, cluster 6 co-expressed chondrocyte markers and cell cycle-related genes (*Mki67*, *Cdk1*, *Stmn1*, *Top2a*, and *Cenpa*), identifying it as a proliferating chondrocyte population (Figure 1A, D). Cluster 1, which expressed mesenchymal and fibroblast markers (*Pdpn*, *Thy1*, and *Clec3b*), was classified as mesenchymal and fibroblastic cells. Cluster 5, characterized by osteoblast markers (*Col1a1*, *Col1a2*, *Bglap*, and *Alpl*), was identified as osteoblasts. Furthermore, cluster 7, which co-expressed fibroblast markers (*Pdpn*) and synovial fibroblast markers (*Prg4*, *Has1*, and *Htra1*), was classified as synovial cells.

Feature plot analysis revealed that *Tmem2* expression was enriched in four specific cell clusters: Chondrocytes (clusters 0, 2, 3, and 4), Proliferating chondrocytes (cluster 6), Osteoblasts (cluster 5), Synovial cells (cluster 7) (Figure 1B, C).

To validate the scRNA-seq findings at the protein level, we next examined TMEM2 expression by immunostaining sagittal knee sections from 14-week-old Tmem2-FLAG knock-in mice. Tmem2 protein was predominantly expressed in articular cartilage chondrocytes, particularly within the non-calcified cartilage layer above the tidemark, with lower expression in both the superficial zone and calcified cartilage (Figure 1E–G). Additionally, Tmem2 protein expression was detected in growth plate chondrocytes, spanning the proliferative to hypertrophic zones (Figure 1H, I). Tmem2 protein expression was also detected in the synovial membrane, specifically in synovial cells known to contribute to hyaluronic acid-rich synovial fluid production. In contrast, no significant expression was detected in the meniscus (Figure 1J, K).

### TMEM2 is transiently upregulated following joint injury

To determine whether the expression pattern of Tmem2 identified in normal mouse knee joints (Figure 1) is altered during osteoarthritis, we next analyzed an independent publicly available scRNA-seq dataset generated from mouse knee joints collected 7 days after non-surgical anterior cruciate ligament (ACL) rupture. The dataset was compared with contralateral intact joints and, as previously reported, was clustered into eight distinct cell populations (Figure 2A).

**Figure 2.**
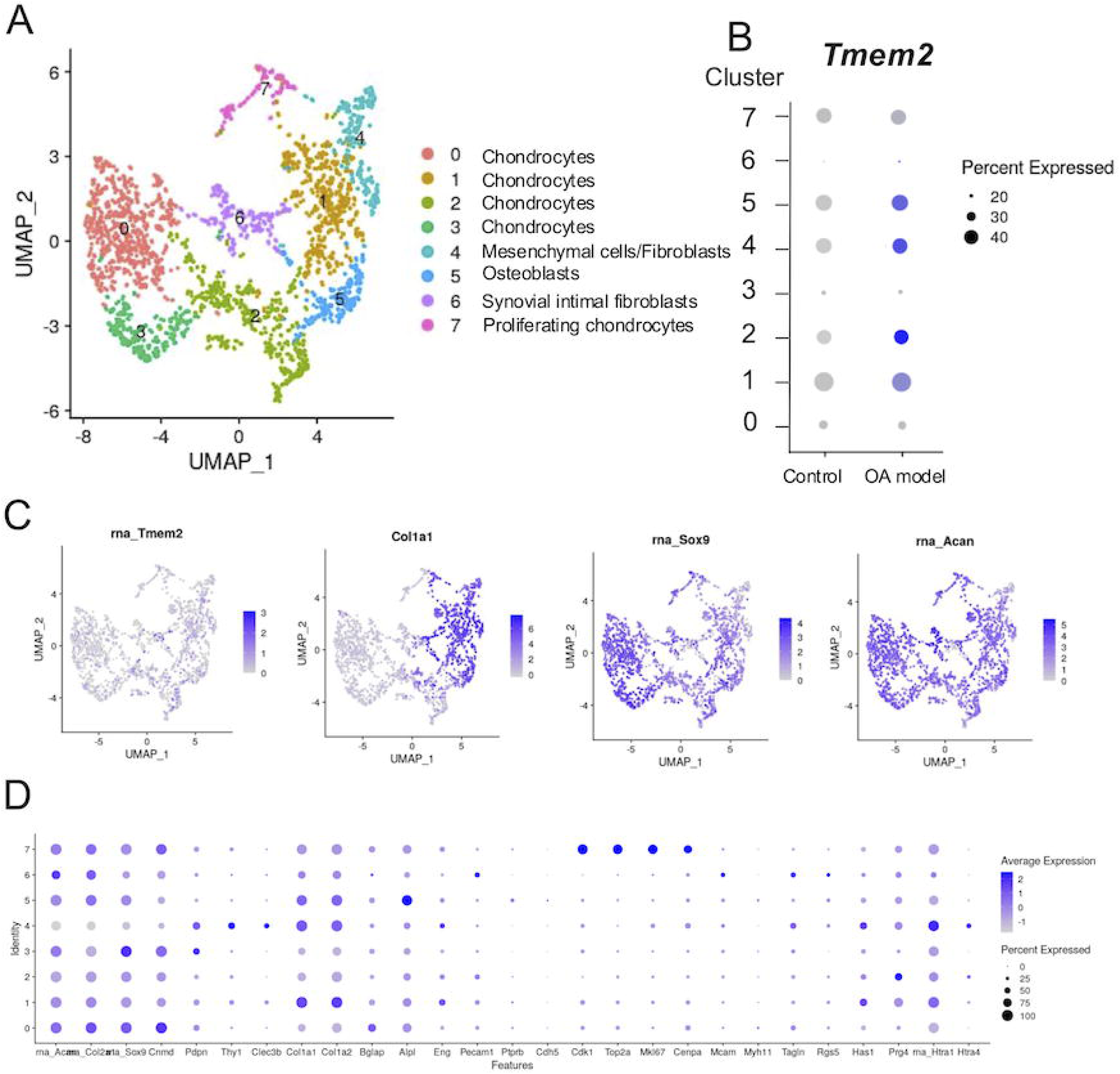
Single-cell RNA-seq analysis of *Tmem2* expression in knee joint chondrocytes post-traumatic osteoarthritis induced by non-surgical ACL rupture. scRNA-seq analysis of osteoarthritic mouse knee joints collected 7 days after ACL rupture. Data are shown for comparison with the normal joint dataset presented in Figure 1. (A) UMAP plot showing eight transcriptionally distinct cell populations identified in mouse knee joint chondrocytes from both control and OA model groups. Clusters are annotated based on known marker gene expression. (B) Dot plot comparing *Tmem2* expression across cell clusters in the control and OA model groups. Increased *Tmem2* expression was observed in clusters 1 and 2 (chondrocytes), cluster 4 (mesenchymal fibroblasts), and cluster 5 (osteoblasts) in the OA model. (C) UMAP feature plots showing expression levels of *Tmem2*, *Col2a1*, *Sox9*, and *Acan* across all clusters. Color scale from grey to blue indicates low to high gene expression intensity. These data demonstrate that the overall cellular composition remained comparable to that of normal joints, whereas TMEM2 expression increased following OA induction. (D) Dot plot presenting representative cell type-specific marker genes used for cluster annotation.

Overall, the major cell populations identified in the OA dataset were comparable to those observed in normal joints, although the relative abundance and transcriptional profiles of several clusters were altered following joint injury.

Clusters expressing chondrocyte markers (*Sox9, Col2a1,* and *Acan*) were annotated as chondrocytes (clusters 0–4). Among these, cluster 2 showed high *Prg4* expression and was annotated as superficial zone chondrocytes (Figure 2A,D). Cluster 7 co-expressed chondrocyte markers and cell cycle–associated genes and was classified as proliferating chondrocytes (Figure 2A,D). Cluster 4, expressing mesenchymal/fibroblast markers (*Pdpn, Thy1,* and *Clec3b*), was annotated as mesenchymal/fibroblastic cells. Cluster 5, expressing osteoblast-associated genes (*Col1a1, Col1a2, Bglap,* and *Alpl*), was annotated as osteoblasts, whereas cluster 6, characterized by *Pdpn, Prg4, Has1,* and *Htra1*, was designated as synovial fibroblasts (Figure 2A,D).

Analysis of the injury-associated dataset revealed increased Tmem2 expression in several cell populations compared with contralateral intact joints, particularly in chondrocytes (clusters 1 and 2), mesenchymal/fibroblastic cells (cluster 4), and osteoblasts (cluster 5), suggesting dynamic regulation of Tmem2 expression following joint injury (Figure 2B).

To determine whether this transcriptional change was also observed in our surgical OA model, we next examined TMEM2 expression in the destabilization of the medial meniscus (DMM) model. In murine DMM models, cartilage degeneration progresses from mild lesions at approximately 2–4 weeks after surgery to more advanced degeneration at later time points, with 4 weeks generally regarded as an early stage of OA progression. Accordingly, we analyzed TMEM2 expression at 4 weeks after DMM to assess early OA-associated changes and compared these findings with those at 8 weeks, representing a more advanced stage of disease.

Immunostaining of sagittal knee sections from the same Tmem2-FLAG knock-in mouse line used in Figure 1 demonstrated detectable TMEM2 expression in articular cartilage in both DMM and naïve joints (Figure 3A–F). However, TMEM2 expression was markedly increased at 4 weeks after DMM, particularly within the non-calcified cartilage layer, and its distribution appeared broader than in naïve controls (Figure 3D,F). In contrast, TMEM2 expression was no longer elevated at 8 weeks after DMM, indicating that its induction was temporally restricted (Supplementary Figure 1). Consistent with these findings, RT-qPCR confirmed increased Tmem2 transcript levels at 4 weeks after DMM, whereas cartilage HA content was significantly reduced (Figure 3G). Together, these findings indicate that transient TMEM2 upregulation during early post-traumatic OA is associated with altered hyaluronan turnover and remodeling of the cartilage extracellular matrix.

**Figure 3.**
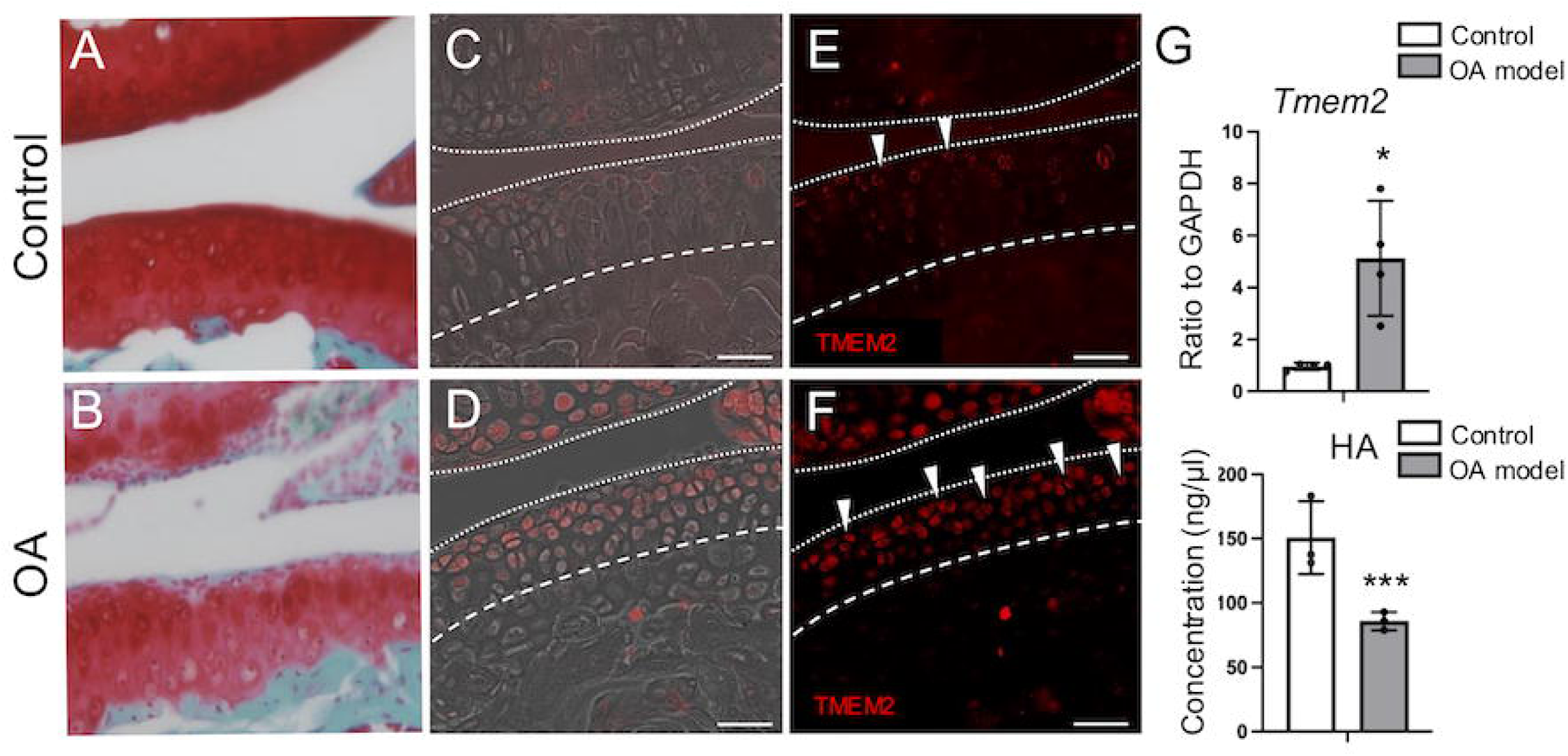
Transient upregulation of TMEM2 during early OA following DMM surgery. Immunostaining was performed using the same Tmem2-FLAG knock-in mouse line shown in Figure 1 following DMM surgery. Representative Safranin O/Fast Green–stained sagittal sections of mouse knee cartilage from control (A) and OA (B) groups. Fluorescence microscopy images of sagittal frozen sections of knee joint articular cartilage obtained 4 weeks after DMM surgery (D, F) or from age-matched naïve controls (C, E). In the OA model, Tmem2 expression was markedly elevated in the non-calcified cartilage layer (arrowheads). (G) Quantitative analysis of Tmem2 and HA expression in articular cartilage. The OA model group showed a significant increase in Tmem2 expression and a significant decrease in HA content compared with controls. Means ± SD (n = 3-4 per group) are shown as horizontal bars. Statistical significance was determined using an unpaired, two-tailed Student’s *t*-test. \***p** < 0.05, \*\*\***p** < 0.001. Scale bars: 50 μm.

Immunostaining results for TMEM2 and HA are shown in Supplementary Figure 2.

### Effect of *Tmem2* Deficiency on Articular Cartilage

To examine whether *Tmem2* deficiency affects chondrocyte homeostasis under physiological conditions, we generated tamoxifen-inducible *Tmem2* conditional knockout (Tmem2-CKO) mice using *Col2a1*-CreERT2. Mice were analyzed at 4, 8, and 12 weeks after tamoxifen administration. Safranin O/Fast Green staining showed that the overall architecture of knee joint cartilage was preserved in both control and Tmem2-CKO mice at all time points examined (Figure 4A–L). However, beginning at 8 weeks after tamoxifen induction, histological analysis revealed an increased number of chondrocytes in Tmem2-CKO mice exhibiting enlarged pericellular spaces and cytoplasmic rarefaction (white arrows, Figure 4F,G). Although such morphological features can occasionally be observed in normal cartilage, they were more frequently detected in Tmem2-CKO mice. These changes are consistent with early degenerative or atrophic-like alterations in chondrocyte morphology (Figure 4A,B,E,F,I,J). No pathological changes were detected in the synovium, subchondral bone, or osteophyte formation (Supplementary Figure 3). Consistent with the established role of TMEM2 in extracellular HA degradation, HA content was significantly increased in articular cartilage from Tmem2-CKO mice compared with controls (Supplementary Figure 4). Thus, loss of Tmem2 alters cartilage HA homeostasis before or in association with the emergence of degenerative morphological changes.

**Figure 4.**
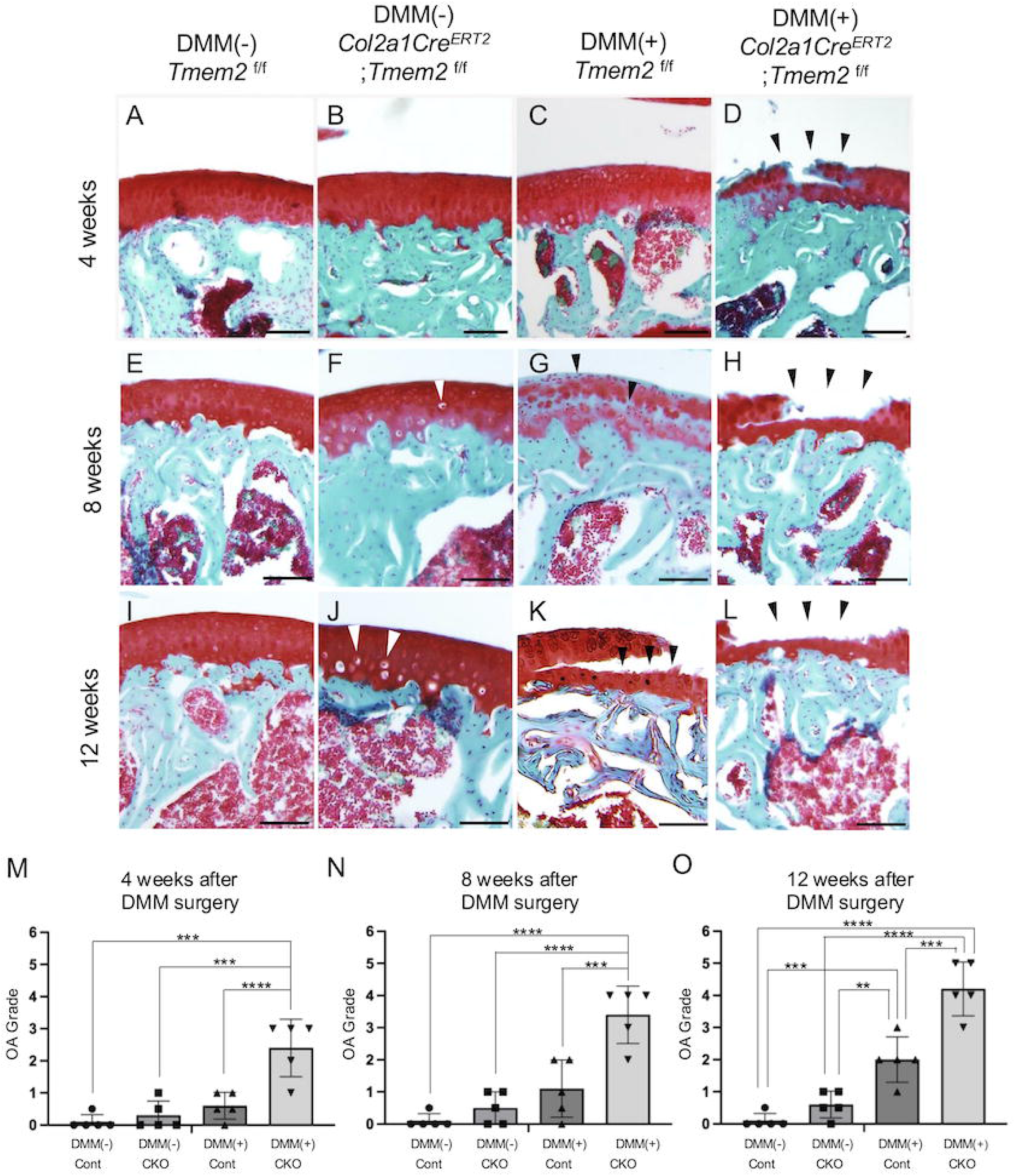
Histological and quantitative analysis of OA progression in *Tmem2*-deficient mice. (A–L) Safranin O/Fast Green–stained sagittal sections of mouse knee cartilage at 4, 8, and 12 weeks after DMM surgery (DMM(+)) or in age-matched naïve mice without DMM surgery (DMM(–)). (F, J) In *Tmem2*-deficient mice (*Col2a1-Cre^ERT2^*; *Tmem2 ^f/f^)* without DMM surgery, Safranin O/Fast Green–stained sections showed chondrocytes with enlarged pericellular spaces and cytoplasmic rarefaction (white arrows), features consistent with early degenerative changes. These cells were more frequently observed in *Tmem2* deficient mice than in control littermates (*Tmem2^f/f^, Tmem2f^/w^*). (D) At 4 weeks post-DMM, *Tmem2*-deficient mice exhibited partial delamination at the interface between non-calcified and calcified cartilage (arrowheads). (G, K) At 8 and 12 weeks post-DMM, control mice showed typical OA features, including reduced Safranin O staining and superficial cartilage fissures (arrowheads). (H, L) In contrast, *Tmem2*-deficient mice exhibited complete cartilage loss at the non-calcified/calcified cartilage interface (arrowheads), indicating markedly accelerated OA progression. (M–O) Quantitative evaluation of OA severity was performed using the Osteoarthritis Research Society International (OARSI) histopathology grading system for mouse cartilage (*n* = 5 mice per group). Each femoral condyle and tibial plateau was graded on a 0–6 scale, and the mean score per joint was calculated from 3–5 serial sections per mouse. No significant differences were observed between control and *Tmem2*-deficient mice in the non-DMM (naïve) group. In contrast, *Tmem2*-deficient mice in the DMM group showed significantly greater OA severity compared to controls. Means ± SD are shown as horizontal bars. *P* values were determined by one-way ANOVA (two-tailed). **p < 0.01, ***p < 0.001, ****p < 0.0001. Scale bars: 100 μm.

### Effect of *Tmem2* Deficiency on OA Pathogenesis and Progression

To further investigate the role of *Tmem2* in OA, we performed destabilization of the medial meniscus (DMM) surgery in Tmem2-deficient mice and littermate controls and monitored disease progression over time. OA severity was evaluated using the Osteoarthritis Research Society International (OARSI) mouse grading system (7-point scale). In control mice, mild OA changes (grade 0.5), characterized by subtle loss of cartilage matrix, were observed at 4 weeks post-DMM (Figure 4C). In age-matched, non-operated Tmem2-deficient mice, focal osteoarthritic features, including chondrocyte nuclear loss, were already evident (Figure 4F,J, white arrows). By 8 weeks post-DMM, OA lesions in control mice progressed to grade 2, with visible matrix degradation and focal delamination of the superficial cartilage layer (Figure 4G, black arrow). At 12 weeks, lesions reached grade 3, showing pronounced erosion of articular cartilage (Figure 4K, black arrow). In contrast, Tmem2-deficient mice subjected to DMM surgery exhibited markedly accelerated OA progression. As early as 4 weeks post-DMM, cartilage delamination at the interface between non-calcified and calcified cartilage was observed, corresponding to grade 3 (Figure 4D, black arrow). By 8 weeks, extensive matrix deterioration resulted in grade 4 OA (Figure 4H), and by 12 weeks, collapse of deep cartilage layers and severe erosion corresponded to grade 5 (Figure 4L). Quantitative histological analysis confirmed significantly more severe cartilage degeneration in Tmem2-deficient mice than in controls (Figure 4M–O). Histopathological assessment of other joint compartments revealed no apparent abnormalities in the synovium, subchondral bone remodeling, or osteophyte formation between genotypes at corresponding time points (Supplementary Figure 3). Together, these findings indicate that the accelerated OA phenotype resulting from Tmem2 deficiency is primarily cartilage-intrinsic and is not secondary to pathological alterations in surrounding joint tissues.

### Effects of *Tmem2* Deficiency on Chondrocyte Survival, Proliferation, and Differentiation

To investigate how *Tmem2* deficiency affects chondrocyte behavior during OA, we performed histological and immunofluorescence analyses. TUNEL staining revealed a significant increase in apoptotic chondrocytes in OA-affected regions of Tmem2-CKO mice (Figure 5A–E). In parallel, the number of Ki-67–positive chondrocytes was markedly reduced (Figure 5F–H), indicating impaired proliferative activity. In contrast, expression of type X collagen was increased (Figure 5I–K), consistent with enhanced hypertrophic differentiation.

**Figure 5.**
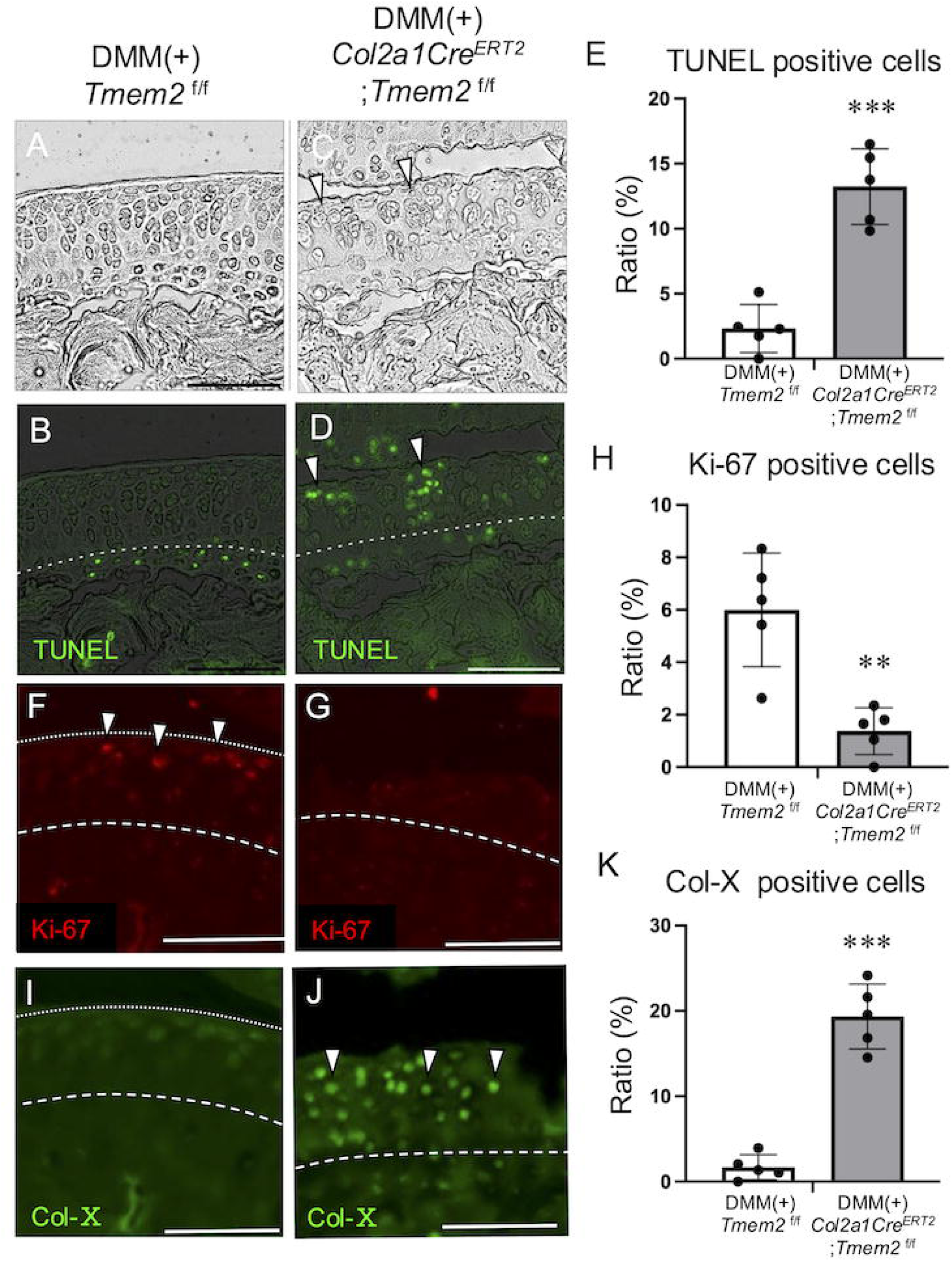
Effects of *Tmem2* deficiency on apoptosis, cell proliferation, and type X collagen expression in OA cartilage. (A–D) Fluorescence microscopy images of sagittal frozen sections of knee joint cartilage 8 weeks after DMM surgery. Apoptotic cells were detected by TUNEL staining, and increased apoptosis was observed in OA-affected regions in *Tmem2*-deficient mice (arrowheads). (E) Quantification of TUNEL-positive cells in the non-calcified cartilage layer. The proportion of TUNEL-positive cells was significantly higher in the OA model group than in the non-OA group in *Tmem2*-deficient mice. (F–G) Immunofluorescence staining for Ki-67 in knee joint cartilage 8 weeks after DMM surgery. Ki-67–positive cells were decreased in OA regions (arrowheads). (H) Quantification of Ki-67–positive cells in the non-calcified cartilage layer. The proportion of Ki-67–positive cells was significantly reduced in the OA model group compared to the non-OA group in *Tmem2*-deficient mice. (I–J) Immunofluorescence staining for type X collagen in knee joint cartilage. Expression was increased in OA regions of *Tmem2*-deficient mice (arrowheads). (K) Quantification of type X collagen–positive cells in the non-calcified cartilage layer. The proportion of type X collagen–positive cells was significantly elevated in the OA model group compared to the non-OA group in *Tmem2*-deficient mice. Means ± SD (*n* = 5) are shown as horizontal bars. *P* values were determined by unpaired, two-tailed Student’s *t*-test. **p < 0.01, ***p < 0.001. This experiment was performed three times with similar results. Data shown in this figure are representative of at least three independent experiments. Scale bars: 100 μm.

Together, these findings indicate that Tmem2 deficiency compromises cartilage homeostasis by promoting chondrocyte apoptosis, reducing proliferative activity, and accelerating hypertrophic differentiation during OA progression.

## Discussion

This study identifies TMEM2 as a critical regulator of HA metabolism in articular cartilage and provides in vivo evidence that TMEM2 contributes to the regulation of hyaluronan turnover within the articular cartilage ECM during osteoarthritis initiation. Under physiological conditions, Tmem2 expression is predominantly localized to the non-calcified zone of articular cartilage. Upon joint destabilization, its expression expands and is transiently upregulated during the early phase of experimental OA. This temporally restricted elevation, supported by single-cell RNA-seq analysis^32^, is consistent with the notion that TMEM2-mediated HA turnover may participate in an early, adaptive response to mechanical or matrix stress.

Consistent with this interpretation, loss of Tmem2 markedly exacerbated OA progression. Conditional deletion of *Tmem2* in chondrocytes resulted in accelerated cartilage delamination, increased apoptosis, reduced proliferative capacity, and enhanced hypertrophic differentiation, indicating that TMEM2 is required to maintain cartilage homeostasis under stress conditions. From an ECM standpoint, HA provides the core scaffold for aggrecan–collagen assemblies that confer compressive resistance and load distribution in cartilage^34,35^. Disruption of HA turnover—either through excessive degradation or impaired clearance—can destabilize pericellular and interterritorial matrix organization, compromise tissue mechanics, and accelerate cartilage degeneration^36^. Our data therefore support a model in which TMEM2 functions as a homeostatic modulator of HA flux, maintaining the synthesis–degradation balance required to preserve matrix organization and mechanics.

Importantly, *Tmem2* deficiency did not produce detectable alterations in synovium, subchondral bone, or osteophyte formation, indicating that the observed degenerative phenotype is primarily cartilage-intrinsic rather than secondary to inflammation or altered subchondral remodeling (Supplementary Figure 3). This cartilage-centric phenotype reinforces TMEM2’s role within the matrix microenvironment that directly surrounds chondrocytes.

At the cellular level, Tmem2 deficiency was associated with increased chondrocyte apoptosis and elevated type X collagen, implicating premature hypertrophic drift. Previous studies have linked TMEM2 loss to apoptosis through mechanisms involving endoplasmic reticulum (ER) stress and dysregulated CD44 signaling ^37,38^. Because mechanical loading can also induce ER stress in chondrocytes^39^, TMEM2-dependent HA remodeling may help stabilize ER homeostasis by maintaining pericellular matrix mechanics and receptor engagement. Defining how TMEM2-controlled HA dynamics intersect with ER stress and survival pathways will clarify the molecular coupling between matrix remodeling, mechanotransduction, and proteostasis in cartilage.

Although the present study establishes an in vivo role for TMEM2 in maintaining cartilage ECM homeostasis, the molecular mechanisms linking TMEM2-dependent HA remodeling to altered chondrocyte survival, proliferation, and differentiation remain to be fully defined. In addition, the upstream signals controlling TMEM2 expression and activity following joint injury are unknown. Further studies will be required to determine how mechanical loading, inflammatory cues, and matrix-derived signals regulate TMEM2 and how altered pericellular HA organization influences receptor signaling, mechanotransduction, and cellular stress responses in chondrocytes.

Collectively, these findings indicate that TMEM2 functions as a context-dependent regulator of cartilage homeostasis. During early OA, transient TMEM2 upregulation may facilitate adaptive ECM remodeling by promoting balanced HA turnover. In contrast, loss of TMEM2 disrupts HA homeostasis, weakens matrix stability, and increases cartilage vulnerability, thereby accelerating degeneration. Together, these observations support a model in which appropriate regulation of TMEM2 activity is required to maintain extracellular matrix integrity, whereas deviation from this balance—particularly through TMEM2 deficiency—predisposes cartilage to degenerative change.

Several limitations of this study should be acknowledged. First, DMM experiments were performed exclusively in female mice. Because female C57BL/6J mice generally develop milder cartilage degeneration following DMM than males, future studies in male mice will be important to determine whether TMEM2 exhibits similar temporal regulation during OA progression irrespective of sex. Second, sham-operated controls were not included because preliminary experiments showed no detectable histological differences from naïve joints. Nevertheless, inclusion of sham-operated controls would further distinguish surgery-related responses from OA-associated alterations. Third, although 4 weeks after DMM is widely regarded as an early stage of OA progression in this model, additional analyses at earlier post-DMM time points would help define the precise onset and duration of TMEM2 induction. Finally, the molecular pathways linking altered TMEM2-dependent HA homeostasis to changes in chondrocyte survival and differentiation remain to be determined.

Although OA-associated TMEM2 mutations have not been reported in humans, the characteristic reduction in HA molecular weight observed in human OA, together with the clinical use of intra-articular HA, highlights the translational relevance of HA metabolic regulation. Rather than broad inhibition or enhancement of HA degradation, these findings suggest that preserving appropriate HA turnover dynamics—potentially through phase-specific modulation of TMEM2 activity—may represent a strategy to maintain ECM architecture and delay OA progression.

## Experimental Procedures

### Animal

All animal experiments followed ARRIVE (Animal Research: Reporting of In Vivo Experiments) guidelines, and a strict protocol was approved by the Institutional Animal Care and Use Committee of Osaka University Graduate School of Dentistry (Approval No. 04263, 04315). A conditional *Tmem2* null allele (*Tmem2^flox^*) and Tmem2-Flag knock-in mice were described previously^26,40^.

*Col2a1Cre^ERT2^* mice (with permission from Prof. Di Chen, Rush University) were generously provided by Prof. Taku Saito, University of Tokyo. We crossed *Col2a1Cre^ERT2^* mice with *Tmem2^f/f^* mice to generate *Col2a1Cre^ERT2^*; *Tmem2 ^f/+^* mice. For breeding, *Col2a1-Cre^ERT2^*; *Tmem2 ^f/+^* and *Tmem2 ^f/f^* mice were used to generate *Col2a1-Cre^ERT2^*; *Tmem2 ^f/f^* mice. Primers used for genotyping are listed in Supplementary tables.

Tamoxifen (Sigma-Aldrich) was dissolved in corn oil (Sigma-Aldrich) to prepare a 10 mg/ml tamoxifen solution. This solution was administered intraperitoneally to 8-week-old *Tmem2 ^f/f^* mice at 200 µl per dose for five consecutive days to generate cartilage-specific *Tmem2* knockout mice^41,42^. All mice were on a C57BL/6J background. Only female mice were used in this study. For DMM surgery, 14-week-old mice were used unless otherwise stated.

For protein localization studies under physiological and OA conditions (Figures 1 and 3), *Tmem2-FLAG* knock-in mice were used. For functional analyses of *Tmem2* deficiency (Figures 4 and 5), tamoxifen-inducible *Col2a1-CreERT2;Tmem2^fl/fl^* mice and littermate controls were used.

### Preparation of paraformaldehyde-fixed cryosections

The knee joints were extracted from the mice and fixed in 4% paraformaldehyde phosphate-buffered saline (PFA) for 24 hours. After fixation, the samples were decalcified by immersion in Osteosoft^®^ (Sigma-Aldrich) at room temperature for one month. After decalcification, the samples were washed three times with 1% bovine serum in phosphate-buffered saline (PBS) and then subjected to sucrose treatment. The samples were embedded in Tissue-Tek (OCT compound; Sakura) as previously described^43^.

### Immunohistochemical staining

Frozen sections of 12 µm thickness were used for immunostaining. Immunostaining of frozen sections was performed as previously described^43^. The primary antibodies used in this study were: rabbit polyclonal ANTI-FLAG (Sigma-Aldrich, F7425, 1:200), biotinylated hyaluronan binding protein (bHABP) was used for detection of HA as previously described^26^. For cell death detection, in situ cell death detection kit, Fluorescein (Roche) was utilized. The detection of proliferating cells was performed using Anti-Ki67 (Abcam, ab16667). For the detection of collagen type X, an anti-collagen type X antibody (Cosmo Bio, LB-0092, 1:200) was used. The secondary antibodies used in this study were: donkey anti-rabbit IgG Alexa Fluor 488(Invitrogen, A32790, 1:500), streptavidin, Alexa Fluor 555 conjugate (Invitrogen, S32355, 1:250) and DAPI (Dojindo, D523, 1:200) from Invitrogen. Mount and coverslip the sections with aqueous fluorescence mounting medium (DAKO). Immunohistochemical evaluation performed by all -in-one fluorescence microscope (Keyence BZ-X710).

As a negative control, tissues with inherently low Tmem2 expression (e.g., spleen and skeletal muscle) were stained under identical conditions. These controls showed no detectable non-specific fluorescence, confirming the specificity and low background of the immunostaining procedure. All staining experiments were repeated at least three times using biologically independent samples to ensure reproducibility.

### RNA Extraction and RT-qPCR Analysis

For RNA analyses, articular cartilage was carefully dissected from both the femoral condyles and tibial plateaus of the knee joint. Prior to tissue collection, all surrounding tissues, including the synovium, menisci, ligaments, joint capsule, and subchondral bone, were completely removed to ensure that only articular cartilage was analyzed. Knee cartilage was extracted from 14-week-old control mice and snap-frozen at -80°C. The cartilage tissue, including the non-calcified articular cartilage layer, was carefully dissolved in TRIzol reagent (Thermo Fisher Scientific) and chloroform (Nacalai Tesque) was added to induce phase separation. The mixture was centrifuged at 13,200rpm for 20 minutes at 4 °C. The aqueous phase was collected and mixed with 70% ethanol, then loaded onto a RNeasy Mini Kit spin column (Qiagen) for RNA purification. The column was centrifuged at 12,000 rpm for 1 minute at room temperature (25 °C), and the flow-through was discarded. Buffer RW1 was added to the column, followed by centrifugation at 12,000 rpm for 30 seconds. After discarding the flow-through, Buffer RPE was added, and the column was centrifuged again at 12,000 rpm for 2 minutes. The flow- through was discarded, and RNase-free water was added to elute the purified RNA.

The extracted RNA was reverse-transcribed into cDNA using oligo(dT) primers and reverse transcriptase (Takara Bio, Japan). For real-time PCR, aliquots of the resulting cDNA were amplified using TaqMan Fast Universal PCR Master Mix (Applied Biosystems, Foster City, CA, USA). Reactions were run on a StepOne Real-Time PCR System with StepOne Software, version 2.1 (Applied Biosystems).

Gene expression levels were normalized to *Gapdh* as an internal control. All reactions were performed in triplicate to ensure reproducibility. The TaqMan probes for *Tmem2* (Mm00459599_m1) and *Gapdh* (Mm99999915_g1) were purchased from Thermo Fisher Scientific.

Cartilage samples from individual mice were processed independently and were not pooled for RNA extraction or subsequent analyses.

### HA Quantification

Articular cartilage was collected from the knee joints of both non-OA and OA model mice. Articular cartilage samples from individual mice were processed and analyzed independently without pooling. Briefly, articular cartilage was carefully dissected, weighed, and homogenized according to the manufacturer’s protocol before HA quantification. The HA content in the tissue was quantified using a Hyaluronic Acid Assay Kit (Cosmo Bio, Japan). After the addition of biotinylated hyaluronic acid-binding protein (Biotin-HABP) and horseradish peroxidase-conjugated avidin (HRP-Avidin), absorbance was measured. A standard curve was generated from known HA concentrations, and the HA content in each sample was calculated accordingly. Three to four biological replicates were analyzed per group.

### Destabilization of the Medial Meniscus (DMM) Surgery for Induction of OA

DMM surgery was performed to induce osteoarthritis (OA) in mice, as it recapitulates the severity and anatomical distribution of cartilage lesions observed in naturally aged mice^44^. All animals were female C57BL/6J mice and surgery was conducted at 14 weeks of age. Mice were anesthetized with an intraperitoneal injection of a triple-agent mixture (medetomidine 0.3 mg/kg, midazolam 4 mg/kg, and butorphanol 5 mg/kg, intraperitoneal). Adequate depth of anesthesia was confirmed by the absence of pedal withdrawal reflex, and body temperature was maintained on a warming pad throughout the procedure. The right knee was shaved and disinfected with povidone–iodine and 70% ethanol. Atipamezole 0.3 mg/kg s.c. was administered at the end of surgery to reverse medetomidine. Pre-emptive analgesia was provided (meloxicam 1 mg/kg s.c.), then continued once daily for 48 h postoperatively. A ∼5 mm vertical skin incision was made from the patella to the tibial tuberosity. The joint capsule was incised medial to the patellar tendon, and the medial meniscotibial ligament (MMTL) was exposed by blunt dissection. Hemostasis was achieved using topical epinephrine. The patella was gently displaced medially to visualize the MMTL, which was then transected with a microsurgical scalpel to destabilize the medial meniscus. The joint capsule was closed with 8- 0 absorbable sutures and the skin with 6-0 nylon.

Mice were allowed to recover on a warming pad and returned to their home cages with ad libitum access to food and water. Ambulation was unrestricted after recovery. Postoperative analgesia (meloxicam 1 mg/kg, subcutaneous) was continued for 48 hours. Animals were monitored at least once daily for wound condition, body weight, and signs of distress; humane endpoints were defined by institutional guidelines.

Age- and sex-matched naïve (non-operated) mice were used as controls. Sham surgery was not performed in this study. However, in preliminary experiments, we confirmed that sham-operated knees (skin and capsule incision without ligament transection) did not exhibit histological or functional differences compared to naïve joints, particularly in terms of cartilage degeneration and OARSI score (data not shown). Based on these findings, naïve joints were deemed an appropriate comparator for evaluating OA development. Mice were randomly assigned to DMM or naïve groups, and all outcome assessments—including histology, immunostaining, and scoring—were performed blinded to group allocation.

### Single-Cell RNA-seq Data Re-Analysis

To investigate the expression and distribution of *Tmem2* in articular chondrocytes under physiological and osteoarthritic (OA) conditions, we reanalyzed publicly available single-cell RNA sequencing (scRNA-seq) datasets derived from mouse knee joints. For normal joint conditions, scRNA-seq data were obtained from non-operated mouse knee joints reported by Sebastian et al. (GEO accession: GSE172500)^32^. For osteoarthritic conditions, we utilized a dataset generated from mice subjected to non-surgical anterior cruciate ligament (ACL) rupture, also reported by Sebastian *et al*. (GSE172500), which captures early post-traumatic, OA-associated transcriptional changes. Although the OA-inducing mechanism in this model differs from that of destabilization of the medial meniscus (DMM) surgery, the ACL rupture model is widely accepted for investigating early, mechanically induced molecular responses in articular chondrocytes. All scRNA-seq data were processed and analyzed using the Seurat R package (version 5.0). Cells with more than twice the median number of detected transcripts were excluded to remove potential doublets, and cells with mitochondrial gene content exceeding 5% were filtered out as low-quality. Following quality control, gene expression data were normalized using global-scaling normalization, and dimensionality reduction was performed with Uniform Manifold Approximation and Projection (UMAP). Clusters expressing immune or hematopoietic lineage markers were excluded from downstream analyses. Cell clustering was performed using the same marker genes reported in the original study to ensure reproducibility of major joint cell populations. Specifically, *Sox9*, *Col2a1*, and *Acan* were used as markers for chondrocytes; *Mki67*, *Cdk1*, and *Top2a* for proliferating cells; *Pdpn*, *Thy1*, and *Clec3b* for fibroblasts/mesenchymal cells; *Col1a1*, *Bglap*, and *Alpl* for osteoblasts; and *Prg4*, *Has1*, and *Htra1* for synovial fibroblasts. Cell-type annotations were cross-validated with those reported in the original publication. The clustering results and annotations were consistent with the original dataset and are shown here to facilitate comparison with our Tmem2 expression analyses. For comparisons between normal and OA datasets, the same analytical pipeline and clustering parameters were applied to ensure consistency across conditions. *Tmem2* expression was visualized using *FeaturePlot* in Seurat (v4.3.0), and differential expression analyses were performed where appropriate. The expression levels and spatial distribution of *Tmem2* were subsequently examined across chondrocyte clusters in both conditions to identify OA-associated alterations.

### Histological Evaluation of *Tmem2*-Deficient OA Model Mice

Knee joints were harvested from *Col2a1Cre^ERT2^*; *Tmem2^f/f^* mice at 4, 8, and 12 weeks following DMM surgery (n ≥ 5 per group). The joints were fixed in 4% paraformaldehyde (PFA) for 24 hours, decalcified in Osteosoft® (Sigma-Aldrich) for approximately one month, cryoprotected in sucrose, and subsequently embedded. Sagittal sections (12 µm thick) were prepared using a cryostat (Leica). Cryosections of 12 μm thickness were used to preserve cartilage architecture and optimize fluorescence immunostaining throughout the full depth of the articular cartilage. Safranin O (0.1%, Sigma-Aldrich) and Fast Green (0.05%, Sigma-Aldrich) staining were performed to evaluate changes in the cartilage matrix and surface integrity between OA and non-OA groups. Stained sections were examined under a stereomicroscope (Olympus).

### Assessment of OA severity

Osteoarthritis severity was graded using the Osteoarthritis Research Society International (OARSI) scoring system, ranging from grade 0 to grade 6^33^ (Supplementary table2).

### Detection of Apoptosis in *Tmem2*-Deficient OA Model

Apoptotic cells in frozen knee joint sections from *Tmem2*-deficient OA model mice were detected using TUNEL staining (TdT-mediated dUTP nick-end labeling). The procedure was performed according to the manufacturer’s protocol provided with the In Situ Cell Death Detection Kit (Roche). Fluorescent signals were visualized using a fluorescence microscope.

### Statistics and Reproducibility

All statistical analyses were performed using GraphPad Prism 8 (GraphPad Software). Differences in Tmem2 expression levels, HA content, and the proportion of TUNEL-positive cells between groups were assessed using two-tailed Student’s t-tests. One-way analysis of variance (ANOVA) followed by post hoc multiple-comparison testing (Tukey’s test) was used to evaluate OA progression in Tmem2-deficient mice. A p value < 0.05 was considered statistically significant. Data are presented as mean ± SD, with the number of biological replicates (n) indicated in each figure legend. Statistical tests were two-sided unless otherwise specified.

No covariates were modeled, and no Bayesian or multilevel analyses were performed. Data were assumed to follow normal distributions, consistent with prior studies of murine OA models. No data were excluded from the analyses, and no statistical methods were used to predetermine sample size; however, sample sizes (n ≥ 5 for DMM groups) were based on previous publications reporting similar effect sizes and are sufficient to detect biologically meaningful differences.

Representative images are shown, and all experiments were independently repeated at least three times with similar results to ensure reproducibility. Each analysis was performed using samples from at least three biological replicates per genotype (n ≥ 3). For histological evaluation after DMM surgery, five or more mice per genotype per time point were included (n ≥ 5). Immunostaining experiments were performed in at least three independent replicates. Mice were randomly assigned to DMM or naïve (non-operated) groups prior to surgery using a random number generator. All histological scoring, immunostaining quantification, and statistical analyses were performed by investigators blinded to group allocation.

## Ethics Statement

All animal experiments were approved by the Institutional Animal Care and Use Committee of Osaka University Graduate School of Dentistry (approval numbers 04263 and 04315) and performed in accordance with the ARRIVE guidelines and institutional regulations for animal welfare. Only female mice were used in this study to minimize variability related to sex hormones. No sex-based comparisons were performed. Sample sizes (n ≥ 5 for DMM groups) were based on previous studies reporting similar effect sizes and are consistent with OARSI guidelines for murine OA models. No data were excluded from the analyses. No wild animals or field-collected samples were used in this study.

## Data availability

All data supporting the findings of this study are available within the paper and its Supplementary Information files. The single-cell RNA-seq dataset analyzed in this study was obtained from the GEO database (accession number GSE172500). All numerical source data underlying the graphs and charts in the main and supplementary figures have been included as Supplementary Data files. Any additional raw data or analysis files generated during this study are available from the corresponding author upon reasonable request.

## Acknowledgments

We thank Ms. Yuki Okamoto and Mayumi Yoshimoto for the excellent care and maintenance of our mouse colony and for valuable assistance in the histological, molecular, and protein work. We would like to express our sincere gratitude to Prof. Di Chen and Prof. Taku Saito for generously providing the *Col2a1Cre^ERT2^* mouse, which was invaluable for our experiments in this study.

## Funding

This work was funded by Grants-in-Aid for Scientific Research (24K02652 and 20H03896 to T.I., 24K23546 to T.M.) from the Japan Society for the Promotion of Science and by the Japan Science and Technology Agency FOREST Program (JPMJFR220J to T.I.).

## Declaration of Interest Statement

The authors declare no competing financial interests.

## Author Contributions

Murotani, contributed to data acquisition, analysis, and interpretation, drafted the manuscript; T. Inubushi, contributed to conception and design, data acquisition, analysis, and interpretation, drafted and critically revised the manuscript; Y. Usami, T. Tomohiro, R. Kani, S. Kusano, S. Hisham, W. Deyang contributed to data acquisition, and analysis; Y. Shiraishi, H. Kurosaka, T. Yamashiro critically revised the manuscript; F. Irie and Y. Yamaguchi contributed to data conception and design, critically revised the manuscript. All authors gave final approval and agreed to be accountable for all aspects of the work.

---

**Supplemental Figure 1.**

Representative Safranin O/Fast Green–stained sagittal sections of mouse knee cartilage at OA Grade 1 (A) and OA Grade 2 (D). Fluorescence microscopy images of sagittal frozen sections of articular cartilage following DMM surgery (B, C, E, F). In the OA Grade 1 group, Tmem2 protein expression was elevated in the non-calcified cartilage layer (B, C), whereas in the OA Grade 2 group, Tmem2 expression was reduced (E, F). White arrows indicate areas with strong Tmem2 immunofluorescence. This experiment was performed using biological replicates (n = 5), and representative images are shown. Data are representative of at least three independent experiments. Scale bars: 20 μm.

**Supplemental Figure 2.**

Fluorescence microscopy images of sagittal frozen sections of mouse articular cartilage 8 weeks after DMM surgery (A–F). In the OA model group, Tmem2 protein expression was elevated in the non-calcified cartilage layer, whereas HA expression was decreased (D–F). White arrows indicate regions of strong HA expression, and yellow arrows indicate regions of strong Tmem2 expression. This experiment was performed using biological replicates (n = 5), and representative images are shown. Data are representative of at least three independent experiments. Scale bars: 20 μm.

**Supplemental Figure 3.**

Representative Safranin O/Fast Green–stained sagittal sections of mouse knee joints from control and Tmem2-deficient (Col2a1-CreERT2; Tmem2^f/f^) mice at 12 weeks post-DMM or from age-matched naïve (non-DMM) mice. In the DMM-operated groups, Tmem2-deficient mice exhibited partial delamination within the articular cartilage (arrowheads). No pathological alterations were detected in the synovium, subchondral bone, or osteophyte formation, supporting the conclusion that the accelerated OA phenotype in Tmem2-deficient mice is primarily cartilage-intrinsic. Scale bars: 200 μm.

**Supplemental Figure 4.**

HA content in articular cartilage. In the *Tmem2*-deficient mice group, the amount of HA was significantly increased. Means ± SD (*n* = 3) are shown as horizontal bars. *P* values were determined by unpaired student-*t* test. **p<0.01.

## Notes

### Competing Interest Statement

The authors have declared no competing interest.

### Summary of Updates

This version has been revised to improve the clarity, rigor, and reproducibility of the study. The Results and Discussion have been updated to clarify the temporal regulation of TMEM2 following joint injury, the effects of chondrocyte-specific Tmem2 deficiency on cartilage degeneration, and the interpretation of TMEM2-dependent hyaluronan homeostasis. The Materials and Methods and figure legends have been expanded to provide additional details regarding experimental design, tissue collection, sample processing, biological replicates, histological evaluation, and quantitative analyses. The limitations of the study, including the use of female mice, the use of naive rather than sham-operated controls, and the need to define earlier temporal and downstream molecular mechanisms, are now more explicitly discussed. Supporting information and associated descriptions have also been revised for clarity and consistency.

## Reference

1. Hunter, D.J. & Bierma-Zeinstra, S. Osteoarthritis. Lancet 393, 1745–1759 (2019).

2. Hunter, D.J., Schofield, D. & Callander, E. The individual and socioeconomic impact of osteoarthritis. Nature reviews. Rheumatology 10, 437–441 (2014).

3. Cross, M. et al. The global burden of hip and knee osteoarthritis: estimates from the global burden of disease 2010 study. Annals of the rheumatic diseases 73, 1323–1330 (2014).

4. Safiri, S. et al. Global, regional and national burden of osteoarthritis 1990-2017: a systematic analysis of the Global Burden of Disease Study 2017. Annals of the rheumatic diseases 79, 819–828 (2020).

5. Woolf, A.D. & Pfleger, B. Burden of major musculoskeletal conditions. Bulletin of the World Health Organization 81, 646–656 (2003).

6. Carr, A.J. Beyond disability: measuring the social and personal consequences of osteoarthritis. Osteoarthritis and cartilage 7, 213–218 (1999).

7. Chen, D. et al. Osteoarthritis: toward a comprehensive understanding of pathological mechanism. Bone research 5(2017).

8. Laurent, T.C. & Fraser, J.R.E. Hyaluronan. FASEB journal 6, 2397–2404 (1992).

9. Papakonstantinou, E., Roth, M. & Karakiulakis, G. Hyaluronic acid: A key molecule in skin aging. Dermato-endocrinology 4, 258–263 (2012).

10. Zhang, W. et al. A decrease in moisture absorption-retention capacity of N-deacetylation of hyaluronic acid. Glycoconjugate journal 30, 577–583 (2013).

11. T, H. et al. Intercellular adhesion molecule-1 mediates the inhibitory effects of hyaluronan on interleukin-1beta-induced matrix metalloproteinase production in rheumatoid synovial fibroblasts via down-regulation of NF-kappaB and p38. Rheumatology (Oxford, England) 45, 824–832 (2006).

12. Nago, M. et al. Hyaluronan modulates cell proliferation and mRNA expression of adhesion-related procollagens and cytokines in glenohumeral synovial/capsular fibroblasts in adhesive capsulitis. Journal of orthopaedic research 28, 726–731 (2010).

13. Webner, D., Huang, Y. & Hummer, C.D. Intraarticular Hyaluronic Acid Preparations for Knee Osteoarthritis: Are Some Better Than Others? Cartilage 13, 1619S–1636S (2021).

14. Altman, R., Manjoo, A., Fierlinger, A., Niazi, F. & Nicholls, M. The mechanism of action for hyaluronic acid treatment in the osteoarthritic knee: a systematic review. BMC musculoskeletal disorders 16, Article number 321 (2015).

15. Matsumoto, K. et al. Conditional inactivation of Has2 reveals a crucial role for hyaluronan in skeletal growth, patterning, chondrocyte maturation and joint formation in the developing limb. Development 136, 2825–2835 (2009).

16. Moffatt, P. et al. Hyaluronan production by means of Has2 gene expression in chondrocytes is essential for long bone development. Developmental dynamics 240, 404–412 (2011).

17. Primorac, D. et al. Knee Osteoarthritis: A Review of Pathogenesis and State-Of-The-Art Non-Operative Therapeutic Considerations. Genes 11, 854 (2020).

18. Stern, R. Hyaluronan metabolism: a major paradox in cancer biology. Pathologie-biologie 53, 372–382 (2005).

19. Stern, R., Kogan, G., Jedrzejas, M.J. & Soltés, L. The many ways to cleave hyaluronan. Biotechnology advances 25, 537–557 (2007).

20. Lepperdinger, G., Strobl, B. & Kreil, G. HYAL2, a human gene expressed in many cells, encodes a lysosomal hyaluronidase with a novel type of specificity. The Journal of biological chemistry 273, 22466–22470 (1998).

21. AM, A., M, S., M, G. & R, S. Purification and characterization of human serum hyaluronidase. Archives of biochemistry and biophysics 305, 434–441 (1993).

22. Martin, D.C. et al. A mouse model of human mucopolysaccharidosis IX exhibits osteoarthritis. Human molecular genetics 17, 1904–1915 (2008).

23. Totong, R. et al. The novel transmembrane protein Tmem2 is essential for coordination of myocardial and endocardial morphogenesis. Development 138, 4199–4205 (2011).

24. Smith, K.A. et al. Transmembrane protein 2 (Tmem2) is required to regionally restrict atrioventricular canal boundary and endocardial cushion development. Development 138, 4193–4198 (2011).

25. Yamamoto, H. et al. A mammalian homolog of the zebrafish transmembrane protein 2 (TMEM2) is the long-sought-after cell-surface hyaluronidase. The Journal of biological chemistry 292, 7304–7313 (2017).

26. Inubushi, T. et al. The cell surface hyaluronidase TMEM2 plays an essential role in mouse neural crest cell development and survival. PLoS genetics 18, e1009765 (2022).

27. Nag, P. et al. Tmem2 Deficiency Leads to Enamel Hypoplasia and Soft Enamel in Mouse. Journal of dental research 102, 1162–1171 (2023).

28. Xue, J. et al. Addition of High Molecular Weight Hyaluronic Acid to Fibroblast-Like Stromal Cells Modulates Endogenous Hyaluronic Acid Metabolism and Enhances Proteolytic Processing and Secretion of Versican. Cells 9, 1681 (2020).

29. Shiozawa, J. et al. Expression and regulation of recently discovered hyaluronidases, HYBID and TMEM2, in chondrocytes from knee osteoarthritic cartilage. Scientific reports 12, 17242 (2022).

30. Sato, S. et al. Human TMEM2 is not a catalytic hyaluronidase, but a regulator of hyaluronan metabolism via HYBID (KIAA1199/CEMIP) and HAS2 expression. The Journal of biological chemistry 299(2023).

31. Narita, T. et al. TMEM2 is a bona fide hyaluronidase possessing intrinsic catalytic activity. The Journal of biological chemistry 299, 105120 (2023).

32. Sebastian, A. et al. Single-Cell RNA-Seq Reveals Transcriptomic Heterogeneity and Post-Traumatic Osteoarthritis-Associated Early Molecular Changes in Mouse Articular Chondrocytes. Cells 10, 1462 (2021).

33. Glasson, S., Chambers, M., Berg, W.V.D. & Little, C. The OARSI histopathology initiative - recommendations for histological assessments of osteoarthritis in the mouse. Osteoarthritis and cartilage 18 Suppl 3, 23 (2010).

34. Watanabe, H., Cheung, S.C., Itano, N., Kimata, K. & Yamada, Y. Identification of hyaluronan-binding domains of aggrecan. The Journal of biological chemistry 272, 28057–28065 (1997).

35. Matsumoto, K. et al. Distinct interaction of versican/PG-M with hyaluronan and link protein. The Journal of biological chemistry 278, 41205–41212 (2003).

36. Schwartz, N.B. & Domowicz, M. Chondrodysplasias due to proteoglycan defects. Glycobiology 12, 57R–68R (2002).

37. Wu, M. et al. Potential mechanism of TMEM2/CD44 in endoplasmic reticulum stress-induced neuronal apoptosis in a rat model of traumatic brain injury. International journal of molecular medicine 52, 119 (2023).

38. Schinzel, R.T., et al. The Hyaluronidase, TMEM2, Promotes ER Homeostasis and Longevity Independent of the UPRER. Cell 179, 1306–1318 (2019).

39. Tsang, K.Y. et al. Surviving endoplasmic reticulum stress is coupled to altered chondrocyte differentiation and function. PLoS biology 5, e44 (2007).

40. Yuki, T. et al. The cell surface hyaluronidase TMEM2 is essential for systemic hyaluronan catabolism and turnover. The Journal of biological chemistry 297, 101281 (2021).

41. Donocoff, R.S., Teteloshvili, N., Chung, H., Shoulson, R. & Creusot, R.J. Optimization of tamoxifen-induced Cre activity and its effect on immune cell populations. Scientific reports 10, 15244 (2020).

42. Laboratory, J. Intraperitoneal Injection of Tamoxifen for Inducible Cre-Driver Lines. (2023).

43. Toshihiro Inubushi, Satoshi Nozawa, Kazu Matsumoto, Fumitoshi Irie & Yamaguchi, Y. Aberrant perichondrial BMP signaling mediates multiple osteochondromagenesis in mice. JCI insight 2, e90049 (2017).

44. Glasson, S.S., Blanchet, T.J. & Morris, E.A. The surgical destabilization of the medial meniscus (DMM) model of osteoarthritis in the 129/SvEv mouse. Osteoarthritis and cartilage 15, 1061–1069 (2007).

