## Supplementary figures and images for "TMEM2-mediated hyaluronan turnover maintains articular cartilage homeostasis during osteoarthritis progression"

### Supplementary Figure 1

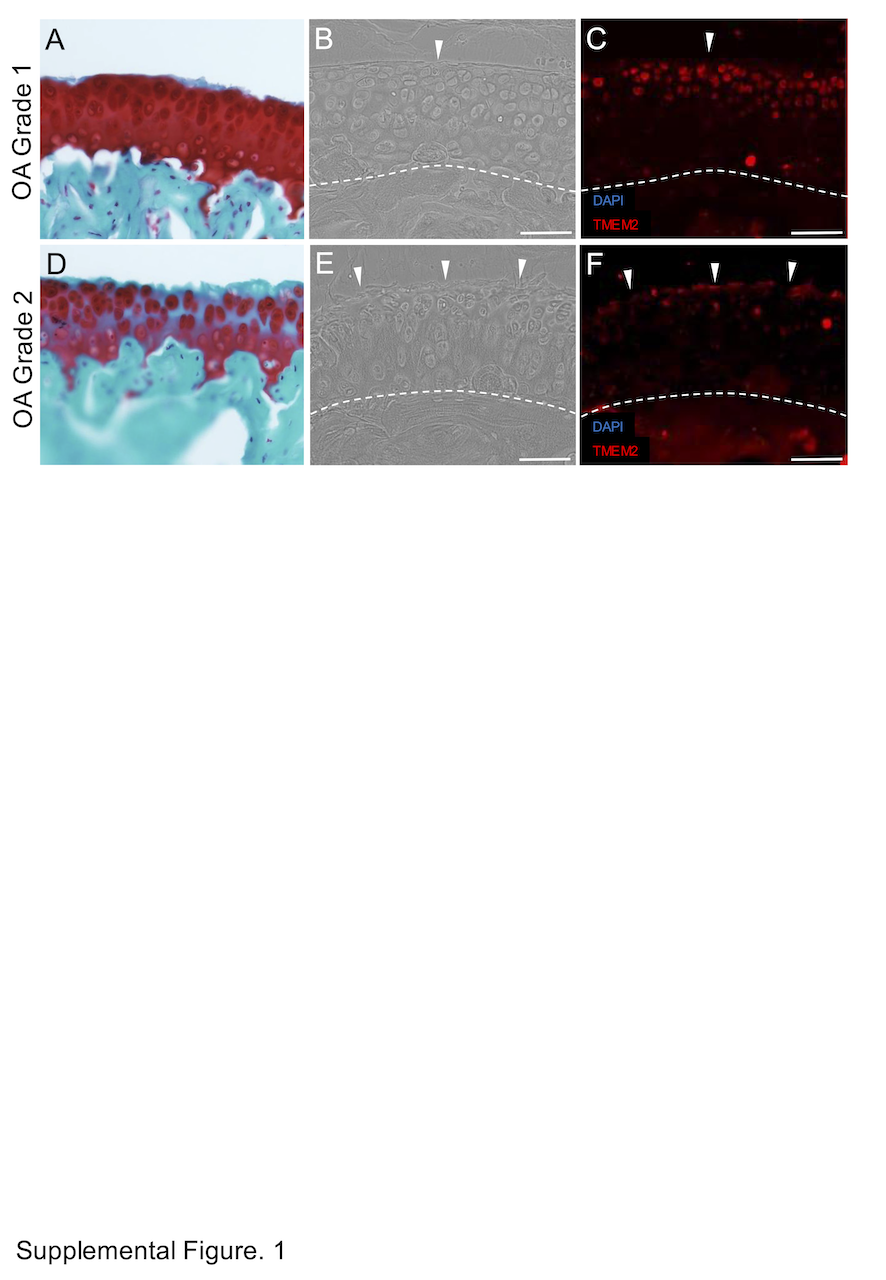

### Supplementary Figure 2

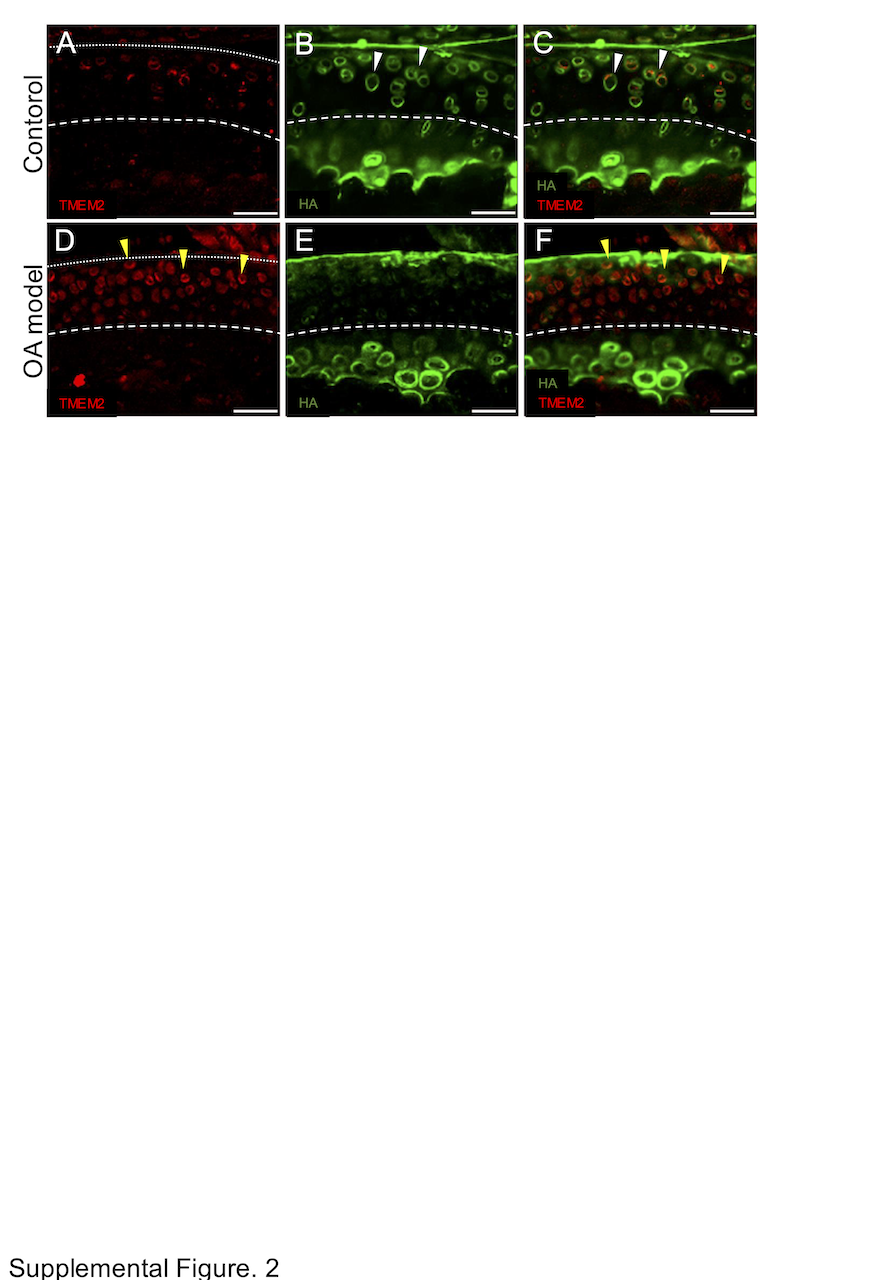

### Supplementary Figure 3

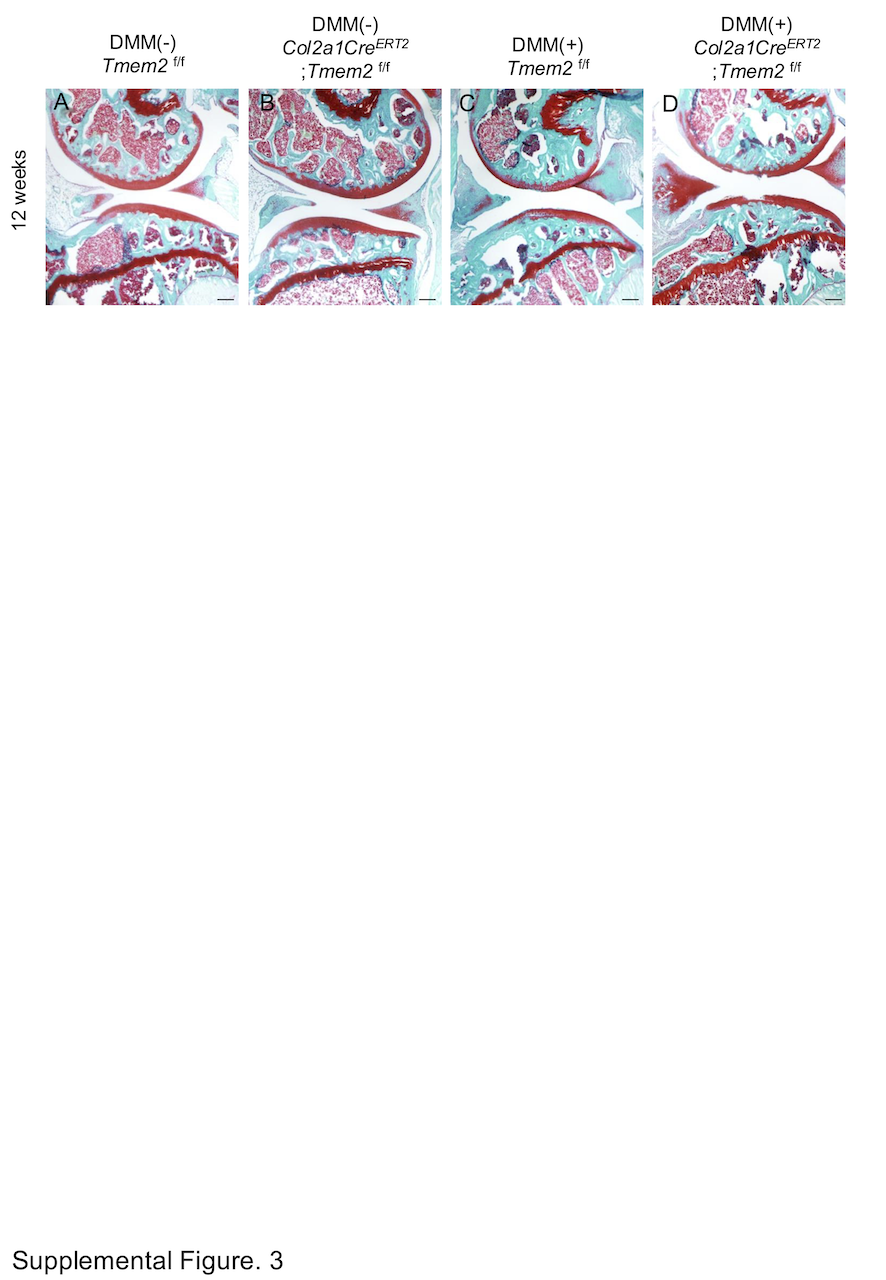

### Supplementary Figure 4

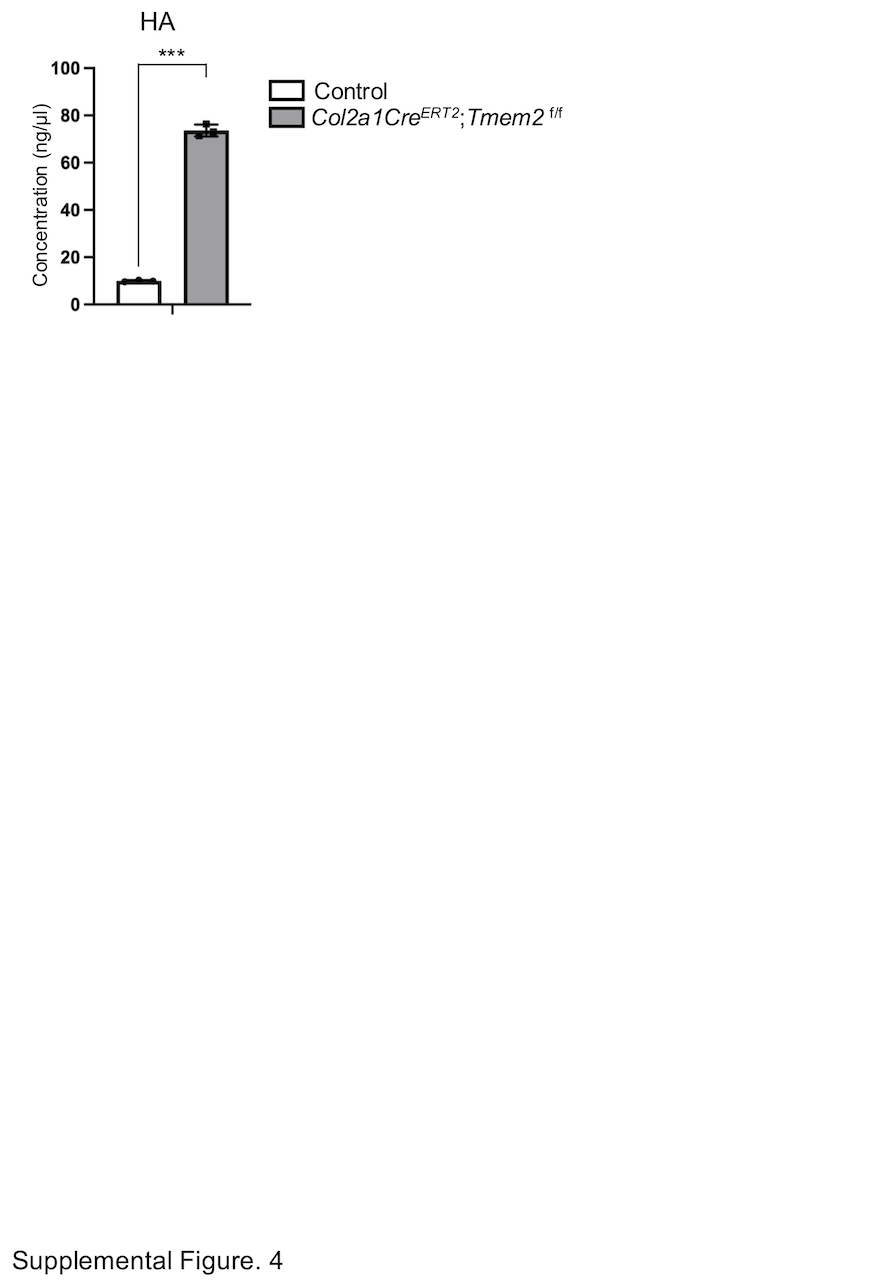
